# Mutations in a novel cadherin gene associated with Bt resistance in *Helicoverpa zea*

**DOI:** 10.1101/698530

**Authors:** Megan L. Fritz, Schyler O. Nunziata, Rong Guo, Bruce E. Tabashnik, Yves Carrière

## Abstract

Transgenic corn and cotton produce crystalline (Cry) proteins derived from the soil bacterium *Bacillus thuringiensis* (Bt) that are toxic to lepidopteran larvae. *Helicoverpa zea,* a key pest of corn and cotton in the U.S., has evolved widespread resistance to these proteins produced in Bt corn and cotton. While the genomic targets of Cry selection and the mutations that produce resistant phenotypes are known in other lepidopteran species, little is known about how Cry proteins shape the genome of *H. zea*. We scanned the genomes of Cry1Ac-selected and unselected *H. zea* lines, and identified eleven genes on six scaffolds that showed evidence of selection by Cry1Ac, including *cadherin-86C* (*cad-86C*), a gene from a family that is involved in Cry1A resistance in other lepidopterans. Although this gene was expressed in the *H. zea* larval midgut, the protein it encodes has only 17 to 22% identity with cadherin proteins from other species previously reported to be involved in Bt resistance. An analysis of midgut-expressed cDNAs showed significant between-line differences in the frequencies of putative nonsynonymous substitutions (both SNPs and indels). Our results indicate that *cad-86C* is a target of Cry1Ac selection in *H. zea*. Future work should investigate phenotypic effects of these nonsynonymous substitutions and their impact on phenotypic resistance in field populations.

## Introduction

Transgenic crops are extensively used for the management of both insect and plant pests worldwide, which places extraordinary pressure on pest species to adapt (Tabashnik and Carrière, 2017; Gould et al. 2018). The first generation of commercially available transgenic crops included corn and cotton bioengineered to produce a single crystalline (Cry) protein from the soil-dwelling bacterium, *Bacillus thuringiensis* (Bt) (Gould, 1998). The primary targets of these Cry proteins were difficult-to-manage lepidopteran larvae (Roush, 1997). When fed upon Cry-producing plant tissue, larvae often experience significant reductions in growth and survivorship (Clark et al., 2000; Tabashnik et al., 2000; Ali et al., 2006; Liu et al., 2010; Pardo-Lopez et al., 2013). Not all targeted lepidopteran species are highly susceptible to Cry proteins, however. Species with low inherent susceptibility are expected (Gould, 1998; Tabashnik et al., 2004; Carrière et al., 2015) and do evolve resistance to Bt crops faster than species with high susceptibility (Tabashnik and Carrière, 2017). Accordingly, there is growing concern over the number of species that have evolved significant resistance to Bt crops producing Cry proteins (Tabashnik and Carrière, 2017; Gould et al., 2018; USEPA, 2018; Tabashnik and Carrière, *in press*).

*Helicoverpa zea* is one lepidopteran species targeted by Bt crops with low susceptibility to Cry proteins (Stone and Sims, 1993; Ali et al., 2006; Sivasupramaniam et al., 2008). A major pest of both corn and cotton in the U.S., *H. zea* has evolved resistance to several Cry proteins (Welch et al., 2015; Dively et al., 2016; Reisig et al., 2018; Kaur et al., 2019; Yang et al., 2019). While some efforts have been made to understand the molecular mechanisms underlying Cry resistance in *H. zea* (Caccia et al., 2012; Zhang et al., 2019), we still know little about how selection by Cry proteins has shaped genotype frequencies in this important pest.

Work in other Cry-resistant Lepidoptera provides clues as to which genes are potential targets of selection in *H. zea*, however. When a larva ingests Cry proteins, these toxins bind to receptors in the midgut, which lead to pore formation and lethal midgut cell death (Pardo-Lopez et al., 2013). Disruption of toxin binding to larval midgut receptors is the most common mechanism of resistance (Peterson et al., 2017). Mutations that alter the coding sequence or reduce expression of Cry1A-binding cadherin proteins are associated with resistance to Cry1A toxins in several major lepidopteran pests (Gahan et al., 2001; Morin et al., 2003; Xu et al., 2005; Fabrick et al., 2014; Zhang et al., 2017; Wang et al., 2018).

Here, we used a genome scanning approach to compare previously described Cry1Ac-selected and unselected lines of *H. zea* (Brévault et al., 2013, 2015; Orpet et al., 2015a, 2015b; Welch et al., 2015; Carrière et al., 2018a). We identified six regions of the genome showing signatures of selection, one of which includes a novel gene from the cadherin family, which has been shown to comprise genes associated with Cry resistance in other lepidopteran species (Gahan et al., 2001; Fabrick et al., 2014; Zhang et al., 2017). We compared the predicted protein sequence of the novel cadherin with cadherins involved in Bt resistance in other lepidopteran species, and analyzed both the midgut expression patterns and predicted amino acid sequence differences at this gene between the selected and unselected *H. zea* lines. Finally, we discuss the possible functional roles of this gene in field-evolved *H. zea* resistance to Cry proteins.

## Methods

### Insect material

We conducted our screen for candidate genes associated with Bt resistance in two laboratory-reared lines of *H. zea* with differing susceptibility to Bt toxin. The lines were founded with 180 larvae collected in Georgia from Cry1Ab corn in 2008 (Brévault et al., 2013). F_1_ progeny from field-collected individuals gave rise to two lines, GA and GA-R, which were reared on wheat-germ diet and were unexposed to Bt toxins or selected for resistance to Cry1Ac, respectively. From 2008 to 2012, each line was reared with *ca*. 900 individuals per generation. Between 2008 and 2010, GA-R was selected with Cry1Ac nine times as described in Brévault et al. (2013), which yielded 10-fold resistance to Cry1Ac in GA-R relative to GA and significantly higher survival on Cry1Ac and Cry1Ac + Cry2Ab cotton in GA-R than GA (Brévault et al., 2013). From 2010 to 2012, GA-R was selected 10 times with Cry1Ac. In 2012, GA-R was crossed to GA to generate a new GA-R line, and the new GA-R and original GA lines were split into two subsets, each reared with ca. 600 individuals per generation (Orpet et al., 2015a). The two subsets of each line were crossed every second or third generation to generate two new subsets (Orpet et al., 2015a). The new GA-R was selected for seven consecutive generations with Cry1Ac, which yielded 14-fold resistance to Cry1Ac in GA-R relative to GA (Orpet et al., 2015a) and significantly higher survival on Cry1Ac + Cry2Ab cotton in GA-R than GA (Carrière, unpubl. data). Between 2012 and 2016, GA-R was selected 25 times with Cry1Ac, using methods described in Carrière et al. (2018a). Male pupae sampled for genomic analysis were from generations F52 for GA-R (n = 5) and F72 for GA (n = 5) reared in October 2016.

### DNA isolation and whole genome sequencing

We extracted whole genomic DNA from 5 individuals per line using a Qiagen® DNEasy Blood and Tissue Kit (Qiagen, Inc., Valencia, CA, USA), following a modified version of the mouse-tail protocol. Genomic DNA was submitted to the North Carolina State University Genomic Sciences Laboratory, where it was prepared for sequencing using an Illumina Truseq LT library prep kit (Illumina, Inc. San Diego, CA). We individually barcoded DNA samples from each individual, after which all DNA samples were pooled for sequencing. We sequenced the prepared pooled library on an Illumina NextSeq500 at North Carolina State University Genomic Sciences Laboratory using 150 base-pair (bp) paired-end reads.

### Read processing and mapping

Sequences were quality filtered to remove all reads with more than 30% of bases having a quality score below Q20 using NGS QC Toolkit v. 2.3.3 (Patel et al., 2012). Low-quality ends (<Q20) were trimmed from the 3’ end of remaining reads to improve overall alignment quality (Del Fabbro et al., 2013). Remaining filter-trimmed reads were mapped to the *H. zea* reference genome (Pearce et al., 2017) using Bowtie v. 2.3.2 (Langmead et al., 2009) in end-to-end mode using the highest sensitivity preset parameters (--very-sensitive). Alignment files were cleaned to keep only reads in proper pairs with robust mapping quality (MAPQ ≥ 10) using samtools v. 1.5 (Li, 2011), and PCR and optical duplicates were identified and removed using picard v. 2.10.5 (http://broadinstitute.github.io/picard). The cleaned alignment files were used to call SNPs with samtools v. 1.5 using the mpileup function, and SNP and indel genotypes in Variant Call Formatted (VCF) files were generated using BCFtools. The VCF files were filtered prior to population genomic analysis to only include loci that were: 1) genotyped in at least 50% of individuals, 2) were sequenced to a minimum depth of coverage of 5 and maximum of 2.5 times the mean genome-wide coverage, 3) had a minor allele frequency (MAF) of > 0.1, and 4) were biallelic, using VCFtools v. 0.1.15 (Danecek et al., 2011).

### Genetic diversity

Following genotype quality filtering, levels of genetic diversity within GA and GA-R were estimated using various metrics. We estimated observed (H_O_) and expected (H_E_) heterozygosity, the proportion of polymorphic SNPs (P_N_), and average MAF with PLINK v.1.07 (Purcell et al., 2007). Inbreeding coefficients (*F*_IS_) were estimated with VCFtools v.0.1.15. To estimate the proportion of shared alleles within lines we calculated D_ST_ in PLINK, with pairwise genetic distance (D) calculated as D = 1-D_ST_.

### Identifying selective sweeps

To identify putative selective sweeps, we searched for regions of the genome with high divergence between GA and GA-R and exceptionally low genetic variation in GA-R, as measured by heterozygosity. We calculated the heterozygosity of within-population pools of individuals (Hp; Rubin et al., 2010) using 40-kb windows with 20-kb of overlap. Hp was calculated as follows:

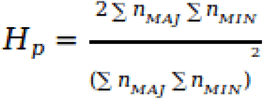

where n_MAJ_ is the number of major alleles and n_MIN_ the number of minor alleles in the window. To prevent spurious signals from few SNPs, we excluded windows with fewer than 10 SNPs. According to Rubin et al. (2010), we standardized estimates of Hp using a Z transformation (ZHp), and putative regions under selection were identified as being over 6 standard deviations from the mean (ZHp < −6). We used *F*_ST_ estimates to identify statistically significant genetic divergence between GA and GA-R. These *F*_ST_ values were calculated two different ways. First, we calculated average *F*_ST_ corresponding to the sliding window averaged ZHp values, using PLINK, and then Z-transformed the average *F*_ST_ values. Next, we calculated sliding window averaged *F*_ST_ values using the p*F*_ST_ function in vcflib (https://github.com/vcflib). *F*_ST_ estimates were calculated over a sliding window size of 5-kb with a 1-kb step, and compared with an empirically-derived estimate of genome-wide divergence between GA and GA-R. A Benjamini and Hochberg (1995) false discovery rate (FDR) correction was applied to the p-values for each of these statistical tests. Regions of the genome with both ZHp < −6 and ZF_ST_ estimates > 1, and an FDR-corrected *F*_ST_ p-value < 0.05 were considered under selection. In the region with strongest evidence of a selective sweep based on the criteria stated above, one of the genes we found was called *cadherin-86C* (*cad-86C*). Because of strong evidence of selection in and around this gene in GA-R, and evidence of cadherin involvement in Bt resistance in other lepidopteran species (Gahan et al., 2001; Morin et al., 2003; Wang et al., 2005; Zhang et al., 2017; Wang et al., 2018), we specifically examined this region further.

### Verification of heterozygosity and genetic divergence at cad-86C by Sanger sequencing

Further evidence for selection at the *cad-86C* gene in GA-R was gathered using a Sanger sequencing approach. Because our whole genome resequencing data were from 5 individuals per line, we amplified and sequenced two ∼600 base-pair (bp) regions of the targeted gene across 24 new individuals in each line to confirm the evidence for selection. These 600 bp target sequences were 1,222 bp apart in a non-coding region of the gene. Primers were designed with PrimerQuest (www.idtdna.com/Primerquest) and are listed in Table S1. Twenty microliter polymerase chain reactions (PCRs) were conducted using 300 ng of genomic DNA per individual, 0.7uM of each forward and reverse primer, 0.2mM dNTPs, and 4uL 5× GoTaq buffer and 1.25 U GoTaq DNA polymerase (Promega Corporation, Madison, WI, USA) according to the recommended protocol. Cycling conditions included denaturation at 95°C for 3 minutes, followed by 30 cycles of 95°C for 30 seconds, 60°C for 30 seconds, 72°C for 30 seconds and a final elongation step at 72°C for 1 minute on Bio-Rad T100 Thermal Cycler (Bio-Rad Laboratories, Inc. Hercules, CA, USA). PCR products were cleaned using an ExoAP reaction (see Supplemental Methods) prior to submission for sequencing. Sanger sequencing was performed using BigDye Terminator v3.1 chemistry (Applied Biosystems) and SNPs were called using PolyPhred (Nickerson et al., 1997). Sequences were manually edited to remove low quality nucleotide calls at their ends, as well as to incorporate variant SNPs for heterozygotes using PolyPhred output. Trimmed and edited sequences were aligned using Clustal Omega available through the European Bioinformatics Institute (EMBL-EBI), and estimates of within population genetic diversity (π, θ_W_) were calculated using the pegas package (v. 0.11; Paradis, 2010) for R (v. 3.5.3; R Development Core Team, 2008). A permutation test, which was custom-coded in R, was used to identify statistically significant differences in π for GA and GA-R at each locus.

**Table 1.**
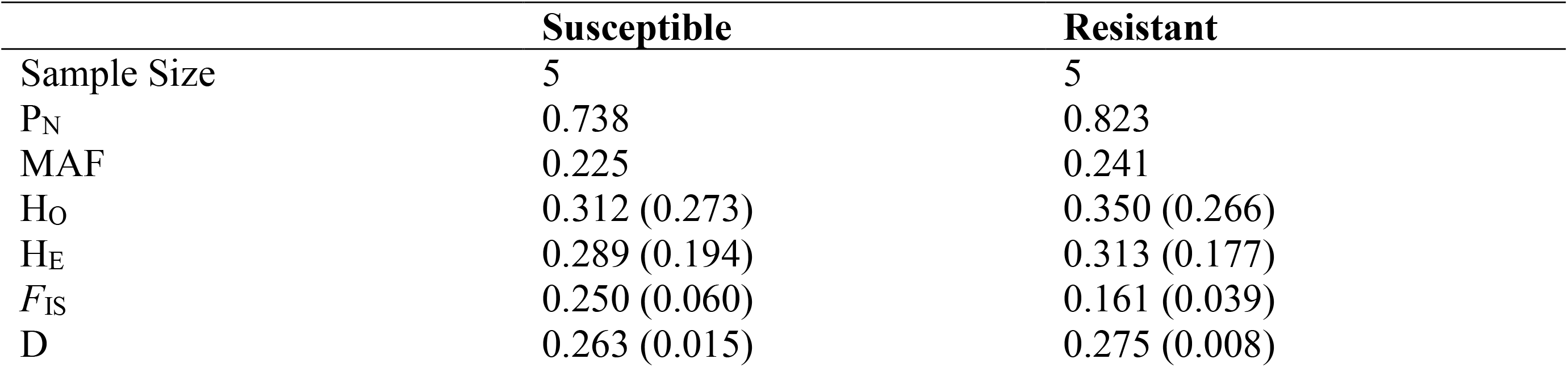
P_N_ is proportion of polymorphic SNPs, MAF is the mean minor allele frequency, H_O_ and H_E_ are observed and expected heterozygosity, and *F*_IS_ is the inbreeding coefficient, and D is the average pairwise genetic distance between individuals in a population. Standard deviations are presented in parentheses.

### Comparison of *CAD-86C* to other cadherin proteins by sequence alignment

We compared our *H. zea* CAD-86C protein sequence (ID = 537580 from the *H. zea* assembly annotation) with CAD-86C orthologues from other species, as well as other cadherins (BtR and CAD2) known to be involved in Bt resistance. All available cadherin protein sequences from 6 lepidopteran species (*H. zea*, *Pectinophora gossypiella, Heliothis virescens, Helicoverpa armigera, Chilo suppressalis, and Bombyx mori*), as well as CAD-86C from *Drosophila melanogaster* were acquired from NCBI. Protein sequences used for analysis, along with their GenBank accession numbers, can be found in Supplementary Data File 1. BtR orthologues and the *C. suppressalis* CAD2 were first aligned to our *H. zea* CAD-86C sequence, and a percentage identity matrix was calculated using T-Coffee available through EMBL-EBI (Madeira et al., 2019). All available BtR, CAD2, and CAD-86C sequences were then aligned to one another using MUSCLE (Edgar et al., 2004), and a phylogenetic analysis was conducted using the phangorn package (v.2.5.3, Schliep, 2011) in R. We used a maximum likelihood approach to distance matrix calculation, assuming a Whelan and Goldman model of molecular protein evolution (Whelan and Goldman, 2001). An unrooted phylogeny was produced using a neighbor-joining approach, and bootstrap support values for the tree nodes were calculated using 1000 resampling events.

### *Cad-86C* expression in the *H. zea* midgut

We reasoned that *cad-86C* should be expressed in the larval midgut to be involved in Bt resistance. To verify midgut expression, we performed reverse transcriptase quantitative PCR (RT-qPCR) on samples from GA and GA-R. We dissected midguts from the F62 generation of GA-R and F81 generation of GA reared in October 2017. GA-R had been selected for resistance to Cry1Ac six times between the F52 generation used for genomic analysis (see above) and the F62 generation. Resistance to Cry1Ac was verified in dissected GA-R larvae by selecting F62 neonates with a diet overlay bioassay using a concentration of 40 µg/cm2 of diet (Carrière et al., 2018a). Susceptibility to Cry1Ac was also verified in dissected GA larvae by selecting F81 neonates with a diet overlay bioassay using a concentration of 30 µg/cm^2^ of diet. Seven days after these bioassays were initiated, third instar (or larger) GA-R larvae were considered resistant while first instar GA larvae were considered susceptible to Cry1Ac. For GA and GA-R, first and third instar (or larger) larvae were respectively transferred to non-Bt diet and their midguts were dissected upon reaching 4th or 5th instar. Dissections were done in ice cold RNAlater+PBS mixture. Whole midguts were stored individually in RNAlater for total RNA extraction.

Total RNA from dissected midguts was isolated using a Zymo Direct-zol RNA miniprep according to the protocol recommended by the manufacturer. We verified RNA quality and determined RNA concentration on an Experion^TM^ automated electrophoresis station. We synthesized first strand cDNA using RevertAid H minus reverse transcriptase and 1µg of total RNA in a 20µL reaction. Gene specific primers were used to amplify *cad-86C* and an endogenous control gene, α-Tubulin (see Table S2 for primer sequences). We performed 20µL qPCR reactions using 10ng of cDNA, 0.5µM primers, and 10uL PowerUp SYBR Green PCR Master Mix (Applied Biosystems). Cycling conditions included initial incubation at 50°C for 2 minutes, followed by denaturation at 95°C for 2 minutes, and then 40 cycles at 95°C for 15 seconds, 60°C for 1 minute on a 7300 Applied Biosystems Real Time PCR System. All reactions were run in triplicate for the 13 GA-R and 14 GA individuals. We observed a single peak in the dissociation curves for all reactions, and PCR efficiencies > 90% for each primer pair using LinRegPCR v.2017.1 (Ruijter et al., 2009). Gene transcript levels were normalized to α-Tubulin and relative expression was standardized using the gene transcript levels detected in GA individuals. Expression of *cad-86C* in GA-R relative to GA was calculated using 2^-ΔΔCt^.

**Table 2.** Estimates of genetic diversity within the GA and GA-R lines in two non-coding regions of the *cad-86C* gene. N represents the number of chromosomes sampled.

| Primer Pair | Sequence Length | Line | N | $\theta_w$ | $\pi$ |
| --- | --- | --- | --- | --- | --- |
| Cad_1b | 421 | GA | 48 | 4.5 | 0.016 |
|  |  | GA-R | 44 | 4.8 | 0.009 |
| Cad_2b | 520 | GA | 48 | 2.0 | 0.006 |
|  |  | GA-R | 48 | 3.2 | 0.004 |

### Long-read sequencing of *cad-86C* cDNAs

To analyze the expressed gene product and putative protein sequence, we designed barcoded PCR primers in the 5’ and 3’ UTR regions of *cad-86C*, amplified the complete cDNA, and sequenced this cDNA by single molecule sequencing. The midgut cDNAs from 13 GA-R and 14 GA individuals were subjected to high fidelity PCRs using the barcoded primer sequences found in Table S3. Full length cDNAs were amplified using Q5 High Fidelity DNA polymerase in mastermix format (New England Biolabs Inc., Ipswich, MA, USA). The 50µL reactions contained 25ng cDNA, 0.5µM primers, and 25µL of 2× Q5 mastermix, and cDNAs were amplified under the following cycling conditions: initial denaturation at 95°C for 3 minutes, followed by 30 cycles of 95°C for 30 seconds, 60°C for 30 seconds, 72°C for 3:40 min and a final elongation step at 72°C for 1 minute on Bio-Rad T100 Thermal Cycler (Bio-Rad Laboratories, Inc. Hercules, CA, USA). Amplicons were run on a 1% agarose gel and the highest molecular weight fragments (> 5kb) were excised, purified using a Zymoclean Gel DNA recovery kit (Zymo Research, Irvine, CA, USA) following the manufacturer’s protocol, and the quantity of purified cDNA was measured using an Agilent D1000 Screentape System (Agilent Technologies, Inc. Santa Clara, CA, USA). Amplicons were then pooled in equimolar amounts, and sequenced on a single Pacific Biosciences (PacBio) SMRT cell at the North Carolina State University Genomic Sciences Laboratory.

**Table 3.**
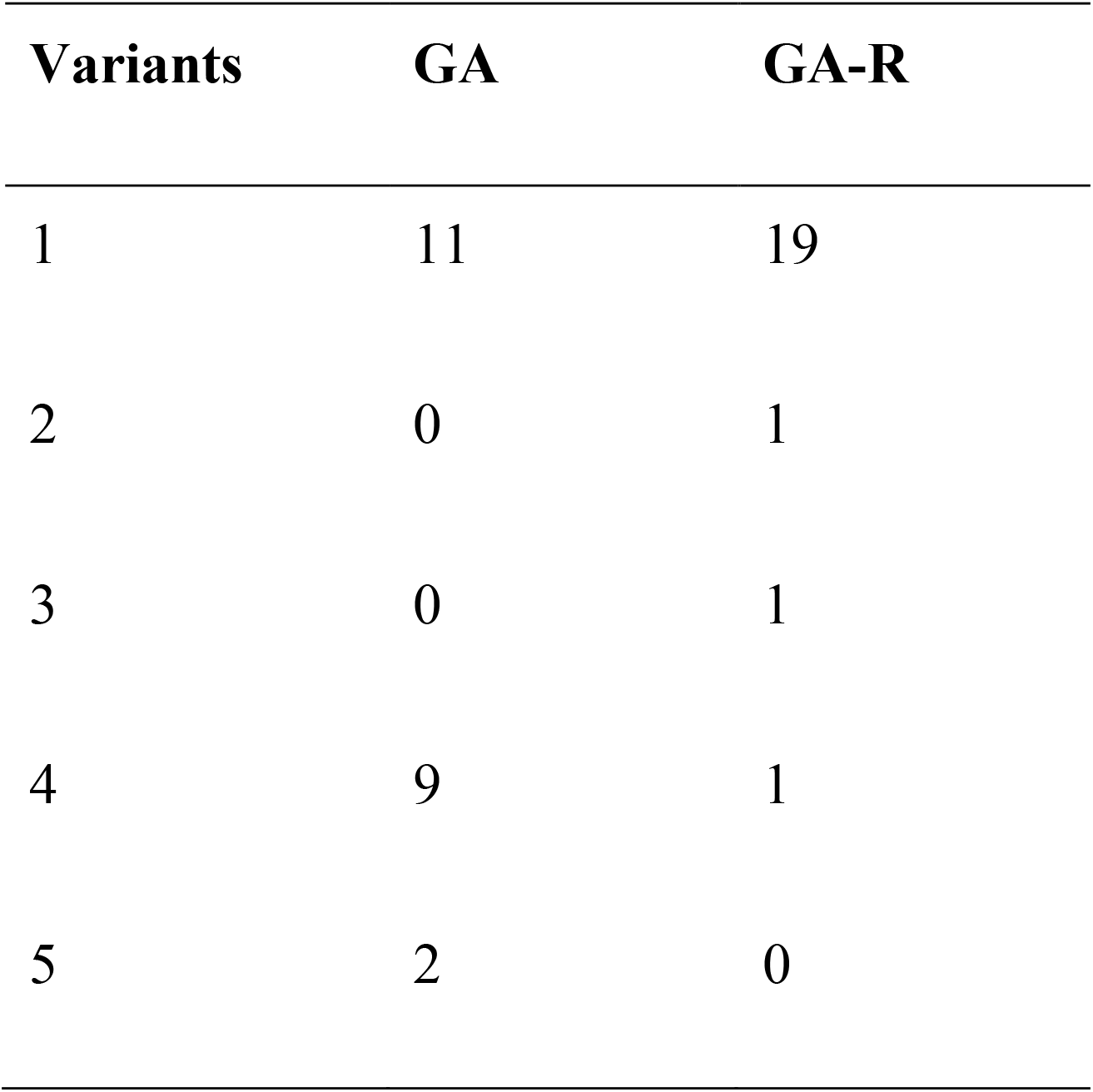
CAD-86C amino acid sequence variant counts for the 22 *H. zea* individuals from the GA and GA-R lines. Counts represent chromosomes sampled (n= 2 per individual) rather than individuals.

### Identification of *CAD-86C* amino acid substitutions

A PacBio-generated bam file was converted to fastq file by bedtools (Quinlan et al., 2010), and sequences from individuals were demultiplexed using the bbduk.sh script from bbmap (Bushnell, 2014). Parameters k and restrictleft were set to 18, which was equal to the primers’ lengths, so that the software only looked for primers matching the leftmost 18 bp. Filtering was done by the FASTA manipulation tool of Galaxy (https://usegalaxy.org; Blankenberg et al., 2010), and the minimal and maximal length parameters were set to 5000 and 0, respectively, to return all sequences longer than 5000 bp. The correction phase of Canu (Koren et al., 2017) was used to improve the accuracy of base calls in PacBio long reads. All error-corrected reads were aligned to the cDNA sequence of *H. zea*’s *cad-86C* gene using the mapPacBio.sh script of bbmap. The results were viewed in Integrative Genomics Viewer (IGV) (Robinson et al., 2011).

Consensus cDNA sequences were identified for each GA and GA-R individual, and ambiguous regions with two alleles, indicating an individual was heterozygous at a locus, were manually edited based upon visual inspection of sequences in IGV. Nucleotide sequences were translated to protein sequences using the translate tool in ExPASy (https://web.expasy.org/translate/), and amino acid sequences were aligned to each other with T-Coffee (Notredame et al., 2000) to identify potential amino acid substitutions. A two-sided Fisher’s exact test was used to compare the distribution of CAD-86C amino acid variants between lines.

## Results

### WGS Data and Variant Call

A total of 427,136,781 raw PE reads were generated, with 356,202,508 remaining after quality filtering, and 179,040,743 remaining after mapping to the reference genome (Table S4). Uniquely placed reads covered 74.3% of the 335.5 megabases (Mb) genome, with a mean genome wide depth of coverage of 16.1× (range 7.96-23.2×) across all individuals. The initial number of variant markers before filtering was 5,106,839. After filtering, the total number of SNPs used for population genomic analysis was 1,986,042, and the total number of short indels was 422,149 (2,408,191 total markers).

### Genetic Diversity and Selective Sweeps

Genome-wide average values of genetic diversity, observed heterozygosity (H_O_), and pairwise genetic distance (D) were similar for GA and GA-R (Table 1). However, GA-R had a lower inbreeding coefficient (*F*_IS_). Overall, the two lines were moderately diverged from each other with a genome-wide *F*_ST_ value of 0.23. Based on heterozygosity estimates among GA-R individuals, a total of 38 genomic windows across 24 scaffolds were identified as putative regions under selection (Figure 1). These candidate regions corresponded to 41 genomic annotations located either upstream or downstream of the identified window (Table S5). The right skewed distribution of ZHp values indicated a high degree of heterozygosity across most 40-kb windows (Figure 1d) throughout the genome. We then examined the Z-transformed *F*_ST_ values across these 40-kb genomic windows, but no 40-kb window had Z*F*_ST_ > 6, which we expected given the moderate genome-wide divergence between lines (Table 1). The maximum Z*F*_ST_ was 3.79 and the Z*F*_ST_ distribution was right skewed (Figure 1c), indicating that there were a number of genomic regions with moderate to high genetic divergence between lines (Figure 1a).

**Figure 1.**
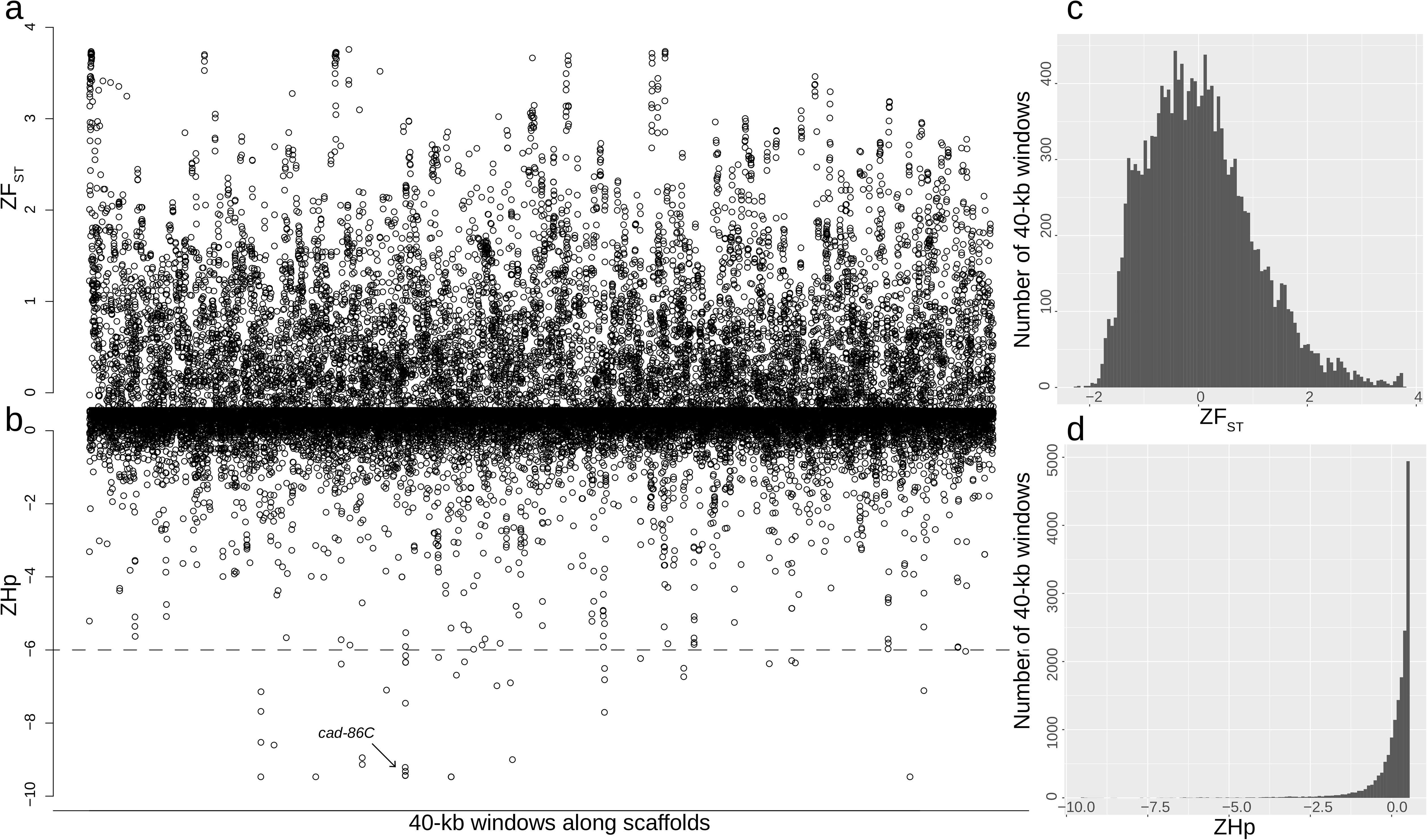
(a) *F*_ST_ between lines and (b) pooled heterozygosity in the GA-R line along 40-kb sliding windows in the *H. zea* genome. The location of the putative *cad-86c* selective sweep is indicated with an arrow in panel b. Panels c and d show the distribution of *F*_ST_ values between lines throughout the genome, and Hp values within the GA-R line throughout the genome, respectively.

When we parsed genomic regions displaying both low Hp (ZHp < −6), high broad-window *F*_ST_ (Z*F*_ST_ > 1), and statistically significant narrow-window *F*_ST_ (p < 0.05), we identified six gene-containing regions with putative selective sweeps (Table S5). These regions were found on six different scaffolds and contained eleven predicted genes: *sol-1* (suppressor of lurcher-1), *cad-86C* (cadherin-86C), *map3k15* (mitogen-activated protein kinase kinase kinase 15), *msp20 (*muscle-specific protein 20), *not1* (CCR4-Not complex subunit 1), *allfused 10*, *cpr47* (cuticular RR1 motif 47), *cgrrf1* (cell growth regulator with RING finger domain 1), *itf46* (intraflagellar transport 46), *cfap100* (cilia and flagellar associated protein 100), and an uncharacterized protein. The *F*_ST_ values for the 40kb regions containing these genes ranged from 0.44 to 0.72, whereas the genome-wide average level of divergence was 0.23. The longest of these putative sweeps extended from 560,000bp on scaffold 20, just upstream of *cad-86C* (ID537580), through the entire length of *cad-86C*, as well as two other genes (an uncharacterized protein and *map3k15*), and ended at bp 740,000. Although the region was broad, the greatest reduction in heterozygosity in GA-R and greatest divergence (*F*_ST_) between lines overlapped with *cad-86C* (Figure 2), a member of a gene family implicated in *Bt* resistance in other lepidopteran species (Gahan et al., 2001; Morin et al., 2003; Wang et al., 2005; Zhang et al., 2017; Wang et al., 2018). This led us to further examine whether *cad-86C* had the potential to serve as target of Cry1Ac selection in *H. zea*.

**Figure 2.**
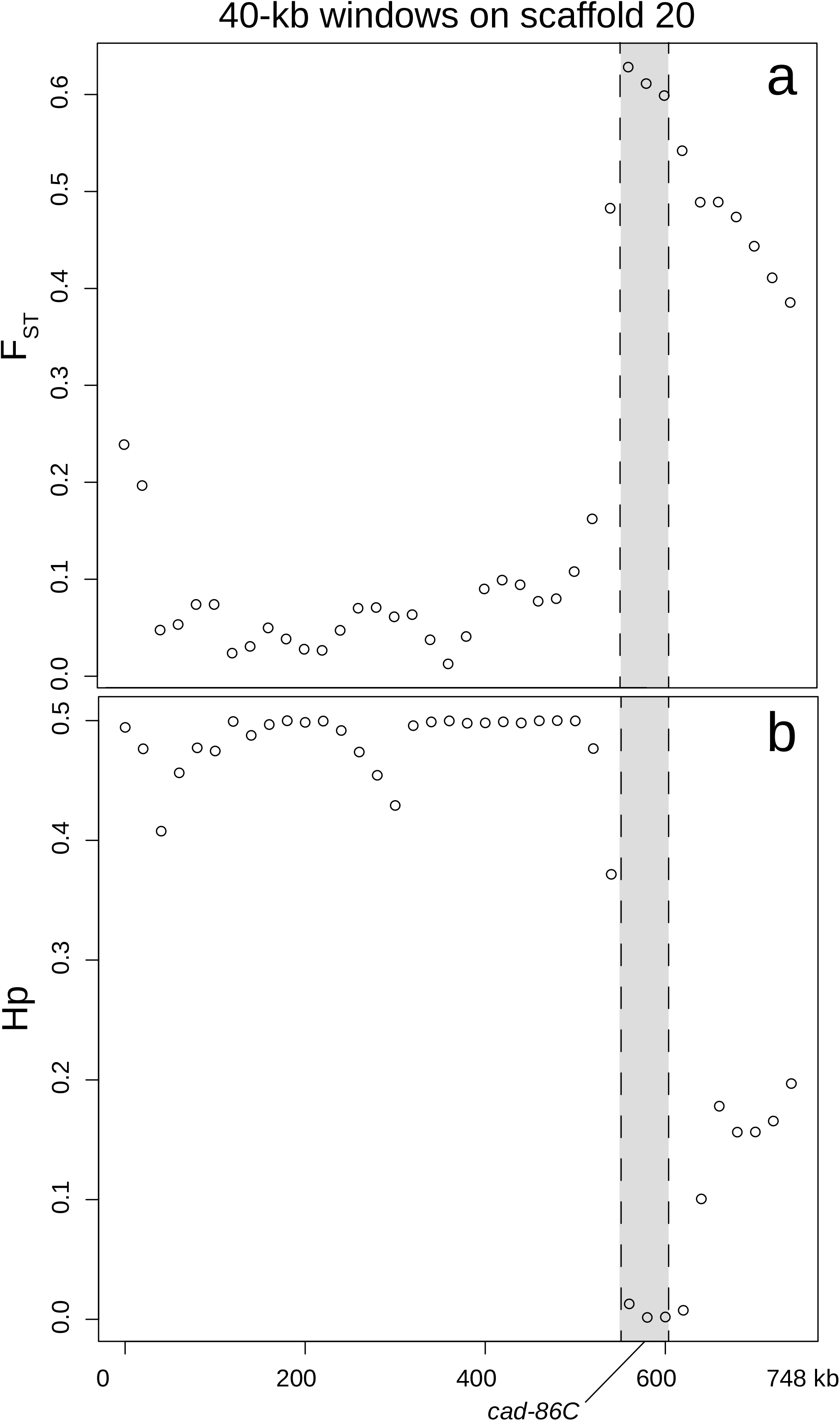
(a) *F*_ST_ between lines and (b) pooled heterozygosity in the GA-R line along 40-kb sliding windows surrounding the putative selective sweep on scaffold 20. *cad-86C* is in grey.

### Verification of heterozygosity and genetic divergence at *cad-86C* by Sanger sequencing

Sanger sequencing of 421 and 520 bp within non-coding regions of *cad-86C* revealed 20 and 9 SNPs in GA and 21 and 14 SNPs in GA-R, respectively (n = 22-24 additional individuals per line). The variation in the numbers of SNPs is reflected in the Watterson’s theta values (θ_W_), which were 4.5 and 2.0 for GA, and 4.8 and 3.2 for GA-R, respectively (Table 2). Within population values of π, however, demonstrated that the allele frequency distributions were different between lines. π was always significantly lower for GA-R than GA according to a permutation test (*p* < 0.001; Table 2; Figures S1 and S2), indicating a reduction in intermediate frequency alleles in GA-R. For the 421 and 520 bp sequences, nucleotide diversity (π) values were 0.016 and 0.006 for GA, and 0.009 and 0.004 for GA-R, respectively.

### Comparison of *CAD-86C* to other cadherin proteins by sequence alignment

Amino acid sequence identity was 56-84% among five cadherin proteins involved in Bt resistance in other Lepidoptera, but only 17-22% between CAD-86C from *H. ze*a and each of these five proteins (Table S6). When we reconstructed the phylogenetic relationships among 14 cadherin proteins, clustering occurred by putative homologues rather than by species (Figure 3). High bootstrap support for the gene clusters demonstrated that CAD-86C was not homologous to other cadherins involved in Bt resistance.

**Figure 3.**
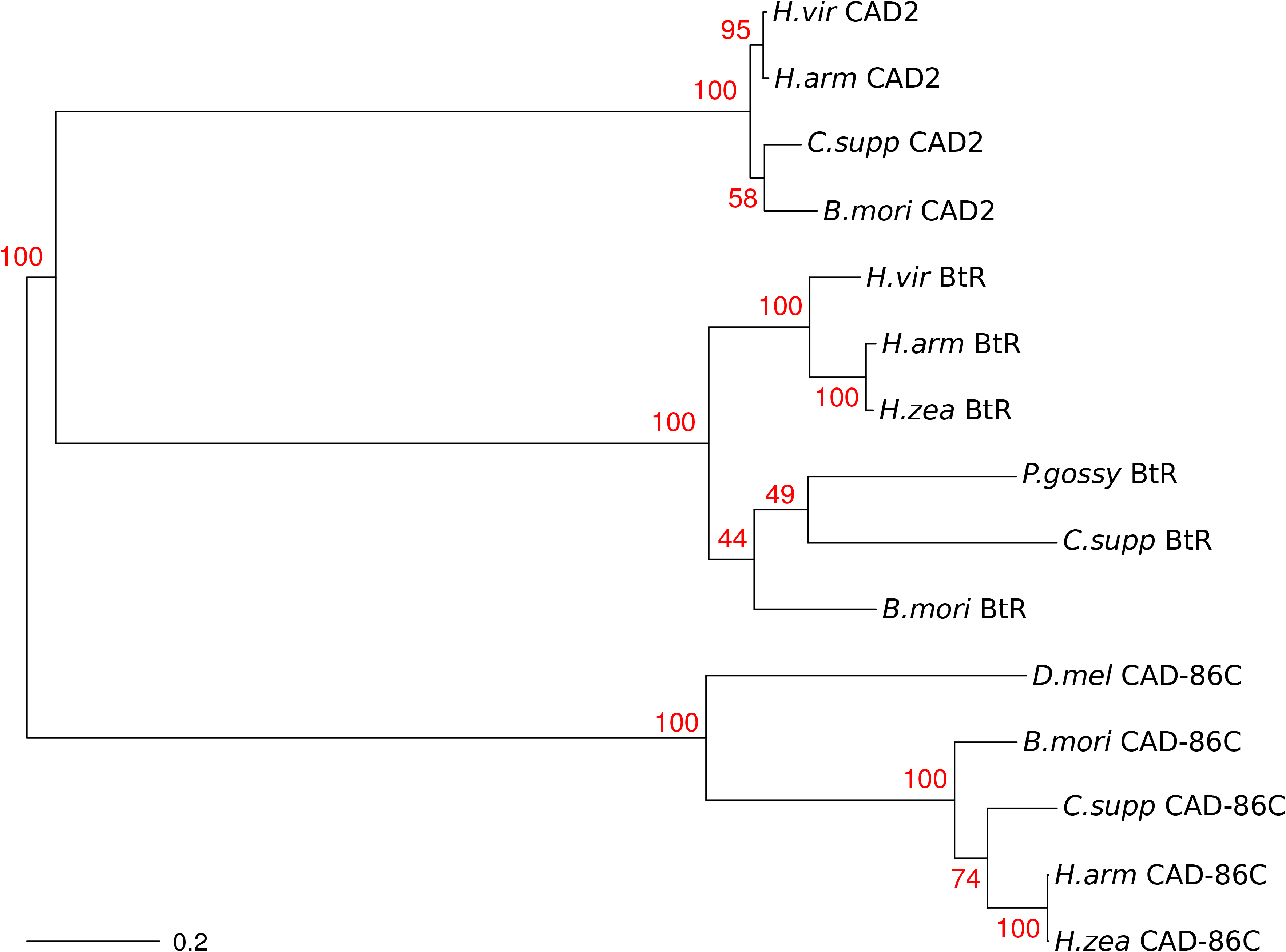
Unrooted neighbor-joining tree indicating the phylogenetic relationships between CAD-86C, CAD2, and BtR. Numbers in red are bootstrap support values (N = 1000) for the tree nodes. A scale bar for genetic distance is in the lower left corner.

### *Cad-86C* expression in *H. zea* midgut

Quantitative PCR revealed that *cad-86C* was transcribed in the midgut of GA and GA-R larvae, with 1.6 fold higher expression in GA-R (Mann-Whitney U-test, *p* = 0.037). Survival to at least 3rd instar was significantly higher for GA-R (61.3%, N = 3840) than GA (1.0%, N = 512) (Fisher’s exact test, *p* < 0.0001) at the time of midgut dissection.

### Identification of *CAD-86C* amino acid substitutions

A 5,539 base pair PCR amplicon, which corresponded to the *cad-86C* coding sequence, was produced for most GA and GA-R individuals (Figure S3). Following PacBio sequencing of these amplicons, a total of 1,081,073 PacBio long reads were generated from 30 *H. zea* individuals. After demultiplexing and quality and length filtering, 14,862 reads remained (minimum length = 5000 bp, Table S7), and 22 individuals (11 per line) were used for analysis.

Using 11 individuals from each line (total n = 22), we identified five predicted CAD-86C protein variants (Figure S4). The distribution of these variant sequences differed significantly between GA and GA-R (Fisher’s exact test, *p* = 0.002; Table 3). Variant 1, the full-length reference protein, was the most common variant in both lines, accounting for 86% of the sequences for GA-R (nine homozygotes and one heterozygote), and 50% for GA (five homozygotes and one heterozygote (Tables 4 and S8). Variants 2 and 3, which occurred in only one individual from GA-R, were identical to the reference sequence except for an insertion at 600,850-600,883 bp of scaffold 20. This insertion introduced a premature stop codon, and was expected to yield a truncated protein. Variant 4, which accounted for 41% of the sequences for GA and 5% for GA-R, had a 15 bp deletion (5’- GAT AAT ACT GCA ACA - 3’) at position 599,832-599,846 bp, and a SNP at 603,408 bp. The deletion was expected to cause the loss of a 5 amino acid sequence (DNTAT) from cadherin repeat domain five, part of the extracellular protein domain which is proximal to the membrane-spanning region. The SNP was expected to produce a threonine to lysine substitution as compared to the reference (T1706K). Variant 5, which appeared in only one individual from GA, had the same 15 bp deletion present in variant 4, and an extra exon in position 600,522-600,574 bp that would terminate the amino acid sequence, producing a truncated protein.

Upon identification of these variants by midgut cDNA sequencing, we revisited our WGS data. All five GA-R individuals were homozygous for the reference sequence. By contrast, only one individual from GA was homozygous for the reference. Two GA individuals were homozygous for the 15 bp deletion leading to the loss of the DNTAT amino acid sequence from the expressed protein, and bore the SNP that caused a threonine to lysine substitution at amino acid position 1706. The final two individuals were heterozygous bearing one copy of the references allele and the alternate allele that produced the DNTAT deletion. This gave a DNTAT + T haplotype frequency of 1 in GA-R, and 0.4 in GA, which was consistent with our PacBio sequencing data.

## Discussion

Here we used whole genome sequencing to identify putative genomic targets of Cry1Ac selection in laboratory-reared *H. zea*, an important agricultural insect pest. This approach has been used previously to identify signatures of selection in other laboratory-selected insects (Izutsu et al., 2012; Jha et al., 2015; Ding et al., 2018). Reduced heterozygosity in an otherwise heterozygous selected population, combined with increased genetic divergence between selected and unselected lines, serves as a signal that a region of the genome was under selection. Because of small population size, genetic drift often occurs among laboratory-reared lines (Schoen et al., 1998; Fritz et al. 2016). When drift occurs, loss of heterozygosity and random fixation of alleles can lead to heightened genome-wide divergence between lines, which can impede separating effects of drift and selection (Hahn, 2019). As recommended by Chandler et al. (2013), we backcrossed GA-R to its ancestral population (GA) and reselected for Cry1Ac resistance, while crossing the two subsets of GA and GA-R every second or third generation to counteract the effects of drift. This led to similar genome-wide observed heterozygosity within each line (0.31 and 0.35 for GA and GA-R, respectively), which was not observed in a pair of laboratory-selected Bt resistant (YHD2) and susceptible (YDK) *H. virescens* lines maintained without backcrossing (Fritz et al., 2016). Furthermore, genome-wide divergence between lines was lower for GA and GA-R (*F*_ST_ = 0.23, n = 5 per line) than YHD2 and YDK (*F*_ST_ = 0.28, n = 43-46 per line). Even so, the genetic divergence between GA and GA-R was slightly higher than that recommended by Perez-Figueroa (*F*_ST_ = 0.2) for a genome scan to identify adaptive loci (Peréz-Figueroa et al., 2010). Therefore, we used both reduced heterozygosity and statistically significant genetic divergence (*F*_ST_) calculated across both broad and narrow sliding genomic windows, to identify genomic regions under selection.

We detected 6 genomic regions with statistically significant evidence of a selective sweep. These genomic regions contained 11 genes, several of which drive various aspects of animal behavior (Table S5). For example, *sol-1* encodes product that is required for proper function of an ionotropic glutamate receptor which regulates synaptic transmission and ultimately locomotory behavior in *Caenorhabditis elegans* (Zheng et al., 2004). *Muscle-specific 20*, another potential locomotory target of selection, is expressed in the muscle tissue of insect larvae and is likely involved in actin binding (Ayme-Southgate et al., 1989). *Map3k15* is a gene from a family involved in the regulation of other genes and ultimately cellular responses to external environmental stimuli (Treisman, 1996). Finally, *cfap* genes are thought to play a role in the motility of cilia, such as those on type-1 sensory neurons in *Drosophila*. Ciliary defects are thought to impact a number of sensory processes (e.g. touch, coordination, taste, olfaction and hearing; Jana et al., 2016).

Previous studies indicate that some species of Lepidoptera show distinct behaviors when exposed to plant tissue or diet treated with Cry proteins as compared to those exposed to untreated tissue or diet. For example, increased locomotory activity and ballooning behaviors are thought to be the result of larval toxin detection and avoidance (Benedict et al., 1992; Men et al., 2005; Prasifka et al., 2009; Goldstein et al., 2010; Ramalho et al., 2014). Furthermore, in binary choice tests, larvae from GA-R preferred to feed on a nutritionally optimal Cry1Ac diet relative to a non-Bt suboptimal diet, while larvae form GA consumed more of the suboptimal non-Bt diet (Orpet et al., 2015b). This suggests that detection and consumption of a nutritionally optimal diet supersedes toxin avoidance behaviors in GA-R. Genes involved in sensory and locomotory function may have served as targets of Cry1Ac selection in GA-R, conferring these behavioral changes. Future work on the function of the genes linked to animal behavior identified here could provide clues as to how behavioral changes occurred in GA-R as a result of Cry1Ac selection.

We focused on a cadherin gene, *cad-86C,* in part because disruption of the coding sequence of cadherin proteins or reduced cadherin gene expression confers resistance to Cry1 toxins in other Lepidoptera (Gahan et al., 2001; Morin et al., 2003; Xu et al., 2005; Fabrick et al., 2014; Zhang et al., 2017; Wang et al., 2018). Moreover, this gene had the second highest level of genetic divergence between lines and second lowest heterozygosity estimates in GA-R in the entire genome (Table S5; Fig. 1). The *H. zea* CAD-86C protein sequence showed only 17-21% identity with other cadherin proteins involved in Bt resistance (this according to Table S6). As far as we know, the function of CAD-86C has been described only for embryonic organogenesis in *D. melanogaster* (Lovegrove et al., 2006; Fung et al., 2008; Schlichting et al., 2008).

We demonstrated that *cad-86C* is transcribed in the midgut of GA and GA-R larvae, which is consistent with a potential role in Bt resistance. The 1.6-fold higher expression of *cad-86C* in GA-R relative to GA was statistically significant, but it seemed unlikely this was the primary cause of the >10-fold difference in resistance between these lines (Orpet et al., 2015a). Therefore, further experiments quantified the difference between lines in the frequency of predicted protein sequence variants. We found that the presence of a 15bp indel, whose *in silico* translation produced amino acid sequence DNTAT, rose from 0.5 in GA to 0.95 in GA-R (Table 3; Figure S4). These frequencies include GA-R individuals with amino acid variants 2 and 3, which also include the 15bp insertion relative to GA variant 4. The same trend was observed in our whole genome sequencing data. The presence of this insertion in the GA line indicated that selection for this mutation in GA-R took place from standing genetic variation in the original field-collected population from Georgia. Interestingly, the mutation was found in the cadherin repeat domain that was most proximal to the membrane-spanning region of the protein. Previous investigations of BtR indicated that the repeat domain most proximal to the membrane-spanning protein domain was critical for toxicity, and mutations in this region promoted Cry1Ab resistance (Hua et al., 2004). A second SNP causing an amino acid substitution appeared to be in strong LD with the 15 bp indel, and the threonine-producing variant rose from 0.55 in GA to 0.95 in GA-R. Putative protein-coding changes, in concert with evidence of midgut gene expression, and the breadth of the putative selective sweep identified by our whole genome sequencing data analysis, provide compelling evidence that this gene is under selection by Cry1Ac in GA-R.

There are a few caveats to interpretation of our findings, however. The first is that identification of a selective sweep at any one of these genes, including *cad-86C*, cannot demonstrate that they are directly involved in Cry1Ac resistance. Instead, it remains possible that some or all of these genomic regions may be under indirect selection. For example, genes in our putative sweep regions may allow for recovery of fitness in individuals bearing mutations with otherwise deleterious effects. Many insecticide resistance mutations are thought to negatively impact fitness in the absence of insecticide pressure (Kliot and Ghanim, 2012), and Bt resistance mutations are no exception (Gassmann et al., 2009; Carrière et al., 2018b). Perhaps selection for a mutation that conferred resistance in GA-R was followed by selection for mutations that improve the fitness of individuals bearing such a mutation.

Secondly, our approach cannot quantify the impact of individual mutations on the trait itself. Rather, by comparing the genomes of multiple populations, it allows us to detect signals that show selection has taken place. It is interesting that the 15bp insertion, which produces a DNTAT amino acid sequence in GA-R, is present in the reference genome. The strain of *H. zea* used for genome sequencing was described as highly susceptible to Cry1A toxins by Pearce et al. (2017), which is true relative to levels of resistance recently described in *H. zea* (Brévault et al., 2013; Dively et al., 2016; Reisig et al., 2018; Yang et al., 2019). Yet in the early 1990s, before Bt crops were commercialized, this long-term laboratory-reared strain was less susceptible to Cry1Ac than most strains derived recently from the field, including a >400-fold difference in one case (Luttrell et al., 1999). Thus, we cannot exclude the possibility that the *cad-86C* allele found here in GA-R and in the strain sequenced by Pearce et al. (2017) decreases susceptibility to Cry1Ac. Additional work remains to characterize the functional role of this and other alleles.

Finally, there are inherent tradeoffs to using laboratory-selected versus field-selected populations to identify genes under selection (Ffrench-Constant, 2013). On the one hand, laboratory-selected populations can provide insights into the genetic mechanisms underlying resistance (Gahan et al., 2001), and Bt resistance mutations identified in laboratory-selected populations have been found in field-selected populations (Zhang et al., 2012; Jin et al., 2018; Mathew et al., 2018; Wang et al., *in press*). On the other hand, some mutations that produce resistant phenotypes in the lab cannot be found in field-collected individuals (Zhang et al., 2012; Fabrick et al., 2014; Mathew et al., 2018). Here, we used a pair of laboratory-reared *H. zea* lines for identification of genes under selection by Cry1Ac. Although we determined that a novel midgut-expressed cadherin gene shows signatures of selection in a Cry1Ac-selected line, it is unclear whether this gene has a significant role in field-evolved Cry1Ac resistance. Future work should examine the importance of *cad-86C* to Bt resistance in field-selected *H. zea*.

## Availability of data and other materials

Whole genome sequencing data and PacBio data are available on the NCBI Sequence Read Archive under accession numbers XXXXXX and XXXXXX (to be updated following acceptance of the publication). Raw Sanger and qPCR data can be found in Dryad Digital Repository XXXXX (to be updated following acceptance of the publication). All scripts used for data analysis can be found at https://github.com/mcadamme/GA_and_GAR.

## Competing interests

The authors declare that they have no competing interests.

## Funding

Funding for this research was provided by the USDA NIFA Biotechnology and Risk Assessment Grant 2016-33522-25640. The funding source did not participate in design of the study and collection, analysis, and interpretation of data and in writing the manuscript.

## Authors’ contributions

YC and BT generated the *H. zea* lines. MLF, SON, YC, and BT designed the experiments. MLF, SON, RG, and YC collected the data. MLF, SON, and RG analyzed the data. MLF, SON, and RG wrote the manuscript. All authors edited and reviewed the final manuscript.

## Acknowledgments

Alex DeYonke and the North Carolina Genomic Sciences Lab prepared samples and sequencing libraries. Anahí Espindola provided valuable feedback on the cadherin phylogeny. Fred Gould read and provided valuable comments on the manuscript.

