## Supplementary material for "Mutations in a novel cadherin gene associated with Bt resistance in *Helicoverpa zea*": FigureS4.pdf

```
T-COFFEE, Version_11.00.d625267 (2016-01-11 15:25:41 - Revision d625267 - Build 507)
Cedric Notredame
CPU TIME:0 sec.
SCORE=1000
*
 BAD AVG GOOD
*
GA_CAD86C_Var1    : 100
GA_CAD86C_Var2    : 100
GA_CAD86C_Var3    : 100
GA_CAD86C_Var4    : 100
GA_CAD86C_Var5    : 100
CAD86C_Ref        : 100
cons              : 100

GA_CAD86C_Var1    MLETLLWPRHTEINRMTFALALVVVWWAVGASAGEPVFDPSTLMRLVLVPADVPVGSVIYRVRASD
GA_CAD86C_Var2    MLETLLWPRHTEINRMTFALALVVVWWAVGASAGEPVFDPSTLMRLVLVPADVPVGSVIYRVRASD
GA_CAD86C_Var3    MLETLLWPRHTEINRMTFALALVVVWWAVGASAGEPVFDPSTLMRLVLVPADVPVGSVIYRVRASD
GA_CAD86C_Var4    MLETLLWPRHTEINRMTFALALVVVWWAVGASAGEPVFDPSTLMRLVLVPADVPVGSVIYRVRASD
GA_CAD86C_Var5    MLETLLWPRHTEINRMTFALALVVVWWAVGASAGEPVFDPSTLMRLVLVPADVPVGSVIYRVRASD
CAD86C_Ref        MLETLLWPRHTEINRMTFALALVVVWWAVGASAGEPVFDPSTLMRLVLVPADVPVGSVIYRVRASD

cons              ******************************************************************


GA_CAD86C_Var1    PDFDYPLHFELIGQMGRLDIGVETLPCTRYNSVCQANIILLRRLEPGRYVDFRLSVRNTRGRSSRI
GA_CAD86C_Var2    PDFDYPLHFELIGQMGRLDIGVETLPCTRYNSVCQANIILLRRLEPGRYVDFRLSVRNTRGRSSRI
GA_CAD86C_Var3    PDFDYPLHFELIGQMGRLDIGVETLPCTRYNSVCQANIILLRRLEPGRYVDFRLSVRNTRGRSSRI
GA_CAD86C_Var4    PDFDYPLHFELIGQMGRLDIGVETLPCTRYNSVCQANIILLRRLEPGRYVDFRLSVRNTRGRSSRI
GA_CAD86C_Var5    PDFDYPLHFELIGQMGRLDIGVETLPCTRYNSVCQANIILLRRLEPGRYVDFRLSVRNTRGRSSRI
CAD86C_Ref        PDFDYPLHFELIGQMGRLDIGVETLPCTRYNSVCQANIILLRRLEPGRYVDFRLSVRNTRGRSSRI

cons              ******************************************************************


GA_CAD86C_Var1    ACSVTGTNATTPRDTIFPHQPGIILVPEDAKRGTDLEIVIARKNPVTPKPLELELWGSPLFAIRQR
GA_CAD86C_Var2    ACSVTGTNATTPRDTIFPHQPGIILVPEDAKRGTDLEIVIARKNPVTPKPLELELWGSPLFAIRQR
GA_CAD86C_Var3    ACSVTGTNATTPRDTIFPHQPGIILVPEDAKRGTDLEIVIARKNPVTPKPLELELWGSPLFAIRQR
GA_CAD86C_Var4    ACSVTGTNATTPRDTIFPHQPGIILVPEDAKRGTDLEIVIARKNPVTPKPLELELWGSPLFAIRQR
GA_CAD86C_Var5    ACSVTGTNATTPRDTIFPHQPGIILVPEDAKRGTDLEIVIARKNPVTPKPLELELWGSPLFAIRQR
CAD86C_Ref        ACSVTGTNATTPRDTIFPHQPGIILVPEDAKRGTDLEIVIARKNPVTPKPLELELWGSPLFAIRQR

cons              ******************************************************************


GA_CAD86C_Var1    RVSAENTEGTIFLVGPLDFEAQSMYHLTLLAVDPFIEIGKDTRNIAGLEVVVVVQDVQDMPPVFTA
GA_CAD86C_Var2    RVSAENTEGTIFLVGPLDFEAQSMYHLTLLAVDPFIEIGKDTRNIAGLEVVVVVQDVQDMPPVFTA
GA_CAD86C_Var3    RVSAENTEGTIFLVGPLDFEAQSMYHLTLLAVDPFIEIGKDTRNIAGLEVVVVVQDVQDMPPVFTA
GA_CAD86C_Var4    RVSAENTEGTIFLVGPLDFEAQSMYHLTLLAVDPFIEIGKDTRNIAGLEVVVVVQDVQDMPPVFTA
GA_CAD86C_Var5    RVSAENTEGTIFLVGPLDFEAQSMYHLTLLAVDPFIEIGKDTRNIAGLEVVVVVQDVQDMPPVFTA
CAD86C_Ref        RVSAENTEGTIFLVGPLDFEAQSMYHLTLLAVDPFIEIGKDTRNIAGLEVVVVVQDVQDMPPVFTA

cons              ******************************************************************


GA_CAD86C_Var1    APPITHLPRQVAPGDMVVRVRAEDGDKGAPRQIRYGLVSEGNPFTPFFNINETSGEVTLERPIEEI
GA_CAD86C_Var2    APPITHLPRQVAPGDMVVRVRAEDGDKGAPRQIRYGLVSEGNPFTPFFNINETSGEVTLERPIEEI
GA_CAD86C_Var3    APPITHLPRQVAPGDMVVRVRAEDGDKGAPRQIRYGLVSEGNPFTPFFNINETSGEVTLERPIEEI
GA_CAD86C_Var4    APPITHLPRQVAPGDMVVRVRAEDGDKGAPRQIRYGLVSEGNPFTPFFNINETSGEVTLERPIEEI
GA_CAD86C_Var5    APPITHLPRQVAPGDMVVRVRAEDGDKGAPRQIRYGLVSEGNPFTPFFNINETSGEVTLERPIEEI
CAD86C_Ref        APPITHLPRQVAPGDMVVRVRAEDGDKGAPRQIRYGLVSEGNPFTPFFNINETSGEVTLERPIEEI

cons              ******************************************************************


GA_CAD86C_Var1    AAISHAGAPILLTVVAEEVRLSREEPEAMSSTVQLAFILPERENSPPYFENQFYITYLDENAPQGT
GA_CAD86C_Var2    AAISHAGAPILLTVVAEEVRLSREEPEAMSSTVQLAFILPERENSPPYFENQFYITYLDENAPQGT
GA_CAD86C_Var3    AAISHAGAPILLTVVAEEVRLSREEPEAMSSTVQLAFILPERENSPPYFENQFYITYLDENAPQGT
GA_CAD86C_Var4    AAISHAGAPILLTVVAEEVRLSREEPEAMSSTVQLAFILPERENSPPYFENQFYITYLDENAPQGT
GA_CAD86C_Var5    AAISHAGAPILLTVVAEEVRLSREEPEAMSSTVQLAFILPERENSPPYFENQFYITYLDENAPQGT
CAD86C_Ref        AAISHAGAPILLTVVAEEVRLSREEPEAMSSTVQLAFILPERENSPPYFENQFYITYLDENAPQGT

cons              ******************************************************************


GA_CAD86C_Var1    ALMFSDPYIPQVNDNDAGKNGVFSLSLVGNNGTFEISPTVAERHAQFIIKVRDNTMLDFEARKSVV
GA_CAD86C_Var2    ALMFSDPYIPQVNDNDAGKNGVFSLSLVGNNGTFEISPTVAERHAQFIIKVRDNTMLDFEARKSVV
GA_CAD86C_Var3    ALMFSDPYIPQVNDNDAGKNGVFSLSLVGNNGTFEISPTVAERHAQFIIKVRDNTMLDFEARKSVV
GA_CAD86C_Var4    ALMFSDPYIPQVNDNDAGKNGVFSLSLVGNNGTFEISPTVAERHAQFIIKVRDNTMLDFEARKSVV
GA_CAD86C_Var5    ALMFSDPYIPQVNDNDAGKNGVFSLSLVGNNGTFEISPTVAERHAQFIIKVRDNTMLDFEARKSVV
CAD86C_Ref        ALMFSDPYIPQVNDNDAGKNGVFSLSLVGNNGTFEISPTVAERHAQFIIKVRDNTMLDFEARKSVV
```

```
cons               ******************************************************************


GA_CAD86C_Var1     FQILAQELGPATNLSVTANVTVYLNDVNDNPPIFLAQSYDVELPENVTAGTKVVQVAADDVDTGAF
GA_CAD86C_Var2     FQILAQELGPATNLSVTANVTVYLNDVNDNPPIFLAQSYDVELPENVTAGTKVVQVAADDVDTGAF
GA_CAD86C_Var3     FQILAQELGPATNLSVTANVTVYLNDVNDNPPIFLAQSYDVELPENVTAGTKVVQVAADDVDTGAF
GA_CAD86C_Var4     FQILAQELGPATNLSVTANVTVYLNDVNDNPPIFLAQSYDVELPENVTAGTKVVQVAADDVDTGAF
GA_CAD86C_Var5     FQILAQELGPATNLSVTANVTVYLNDVNDNPPIFLAQSYDVELPENVTAGTKVVQVAADDVDTGAF
CAD86C_Ref         FQILAQELGPATNLSVTANVTVYLNDVNDNPPIFLAQSYDVELPENVTAGTKVVQVAADDVDTGAF

cons               ******************************************************************


GA_CAD86C_Var1     GKVQFTAILGYLNTSLNLDPISGVITVATNNHGFDREAMPDLHFLVEARDNDGVGLRVTVPLIIKL
GA_CAD86C_Var2     GKVQFTAILGYLNTSLNLDPISGVITVATNNHGFDREAMPDLHFLVEARDNDGVGLRVTVPLIIKL
GA_CAD86C_Var3     GKVQFTAILGYLNTSLNLDPISGVITVATNNHGFDREAMPDLHFLVEARDNDGVGLRVTVPLIIKL
GA_CAD86C_Var4     GKVQFTAILGYLNTSLNLDPISGVITVATNNHGFDREAMPDLHFLVEARDNDGVGLRVTVPLIIKL
GA_CAD86C_Var5     GKVQFTAILGYLNTSLNLDPISGVITVATNNHGFDREAMPDLHFLVEARDNDGVGLRVTVPLIIKL
CAD86C_Ref         GKVQFTAILGYLNTSLNLDPISGVITVATNNHGFDREAMPDLHFLVEARDNDGVGLRVTVPLIIKL

cons               ******************************************************************


GA_CAD86C_Var1     LDVNDNPPEFERSLYEFVLSPSLNNFTSAAFVKAVDRDSEPPNNIVRYEIIQGNGDGKFAINEDTG
GA_CAD86C_Var2     LDVNDNPPEFERSLYEFVLSPSLNNFTSAAFVKAVDRDSEPPNNIVRYEIIQGNGDGKFAINEDTG
GA_CAD86C_Var3     LDVNDNPPEFERSLYEFVLSPSLNNFTSAAFVKAVDRDSEPPNNIVRYEIIQGNGDGKFAINEDTG
GA_CAD86C_Var4     LDVNDNPPEFERSLYEFVLSPSLNNFTSAAFVKAVDRDSEPPNNIVRYEIIQGNGDGKFAINEDTG
GA_CAD86C_Var5     LDVNDNPPEFERSLYEFVLSPSLNNFTSAAFVKAVDRDSEPPNNIVRYEIIQGNGDGKFAINEDTG
CAD86C_Ref         LDVNDNPPEFERSLYEFVLSPSLNNFTSAAFVKAVDRDSEPPNNIVRYEIIQGNGDGKFAINEDTG

cons               ******************************************************************


GA_CAD86C_Var1     ELYLLEALKRTKKQNVHRRRRQSNDQEESEVFVLTIRAYDLGVPRLSSTTIAKVYPPESKTRTMSF
GA_CAD86C_Var2     ELYLLEALKRTKKQNVHRRRRQSNDQEESEVFVLTIRAYDLGVPRLSSTTIAKVYPPESKTRTMSF
GA_CAD86C_Var3     ELYLLEALKRTKKQNVHRRRRQSNDQEESEVFVLTIRAYDLGVPRLSSTTIAKVYPPESKTRTMSF
GA_CAD86C_Var4     ELYLLEALKRTKKQNVHRRRRQSNDQEESEVFVLTIRAYDLGVPRLSSTTIAKVYPPESKTRTMSF
GA_CAD86C_Var5     ELYLLEALKRTKKQNVHRRRRQSNDQEESEVFVLTIRAYDLGVPRLSSTTIAKVYPPESKTRTMSF
CAD86C_Ref         ELYLLEALKRTKKQNVHRRRRQSNDQEESEVFVLTIRAYDLGVPRLSSTTIAKVYPPESKTRTMSF

cons               ******************************************************************


GA_CAD86C_Var1     IVPGANPDKKKLEEVLSTLSGGKVTIIDIKPYRGSDNTATDLNGQESSQEKSEVIAVVRMSGNTAI
GA_CAD86C_Var2     IVPGANPDKKKLEEVLSTLSGGKVTIIDIKPYRGSDNTATDLNGQESSQEKSEVIAVVRMSGNTAI
GA_CAD86C_Var3     IVPGANPDKKKLEEVLSTLSGGKVTIIDIKPYRGSDNTATDLNGQESSQEKSEVIAVVRMSGNTAI
GA_CAD86C_Var4     IVPGANPDKKKLEEVLSTLSGGKVTIIDIKPYRGS-----DLNGQESSQEKSEVIAVVRMSGNTAI
GA_CAD86C_Var5     IVPGANPDKKKLEEVLSTLSGGKVTIIDIKPYRGS-----DLNGQESSQEKSEVIAVVRMSGNTAI
CAD86C_Ref         IVPGANPDKKKLEEVLSTLSGGKVTIIDIKPYRGSDNTATDLNGQESSQEKSEVIAVVRMSGNTAI

cons               ***********************************     **************************


GA_CAD86C_Var1     NVAKLQEQLAKNVTIYTTGTNSGGVINTGTGSGSNSGKSDHASTADGSDSGLYRAESRLLFWLLIL
GA_CAD86C_Var2     NVAKLQEQLAKNVTIYTTGTNSGGVINTGTGSGSNSGKSDHASTADGSDSGLYRAESRLLFWLLIL
GA_CAD86C_Var3     NVAKLQEQLAKNVTIYTTGTNSGGVINTGTGSGSNSGKSDHASTADGSDSGLYRAESRLLFWLLIL
GA_CAD86C_Var4     NVAKLQEQLAKNVTIYTTGTNSGGVINTGTGSGSNSGKSDHASTADGSDSGLYRAESRLLFWLLIL
GA_CAD86C_Var5     NVAKLQEQLAKNVTIYTTGTNSGGVINTGTGSGSNSGKSDHASTADGSDSGLYRAESRLLFWLLIL
CAD86C_Ref         NVAKLQEQLAKNVTIYTTGTNSGGVINTGTGSGSNSGKSDHASTADGSDSGLYRAESRLLFWLLIL

cons               ******************************************************************


GA_CAD86C_Var1     LAILVALVLLLLICCCICEGCPLYMPPRKRVIRVNSTEDDVHLVVRDKGLGRENKSTQNIENKSIQ
GA_CAD86C_Var2     LAILVALVLLLLICCCICEGCPLYMPPRLKCTNM--------------------------------
GA_CAD86C_Var3     LAILVALVLLLLICCCICEGCPLYMPPRLKCTN---------------------------------
GA_CAD86C_Var4     LAILVALVLLLLICCCICEGCPLYMPPRKRVIRVNSTEDDVHLVVRDKGLGRENKSTQNIENKSIQ
GA_CAD86C_Var5     LAILVALVLLLLICCCICEGCPLYMPPRLKCTN---------------------------------
CAD86C_Ref         LAILVALVLLLLICCCICEGCPLYMPPRKRVIRVNSTEDDVHLVVRDKGLGRENKSTQNIENKSIQ

cons               **************************** :  .


GA_CAD86C_Var1     ANEWRRREAWSAEQADLRTKPTQWKFNKRNHRSNKEPSKPASTPGDIHQEFVHTAADNDYRYNDNR
GA_CAD86C_Var2     ------------------------------------------------------------------
GA_CAD86C_Var3     ------------------------------------------------------------------
GA_CAD86C_Var4     ANEWRRREAWSAEQADLRTKPTQWKFNKRNHRSNKEPSKPASTPGDIHQEFVHTAADNDYRYNDNR
GA_CAD86C_Var5     ------------------------------------------------------------------
CAD86C_Ref         ANEWRRREAWSAEQADLRTKPTQWKFNKRNHRSNKEPSKPASTPGDIHQEFVHTAADNDYRYNDNR
```

```
cons

GA_CAD86C_Var1    QSFRRDGPNIIYTKEMQLQEAFANKHKEYIEDLENGYDRIATLHYQRREQDNDSIRRHEIDRGSEV
GA_CAD86C_Var2    ----------------QLQEAFANKHKEYIEDLENGYDRIATLHYQRREQDNDSIRRHEIDRGSEV
GA_CAD86C_Var3    ------------------------------------------------------------------
GA_CAD86C_Var4    QSFRRDGPNIIYTKEMQLQEAFANKHKEYIEDLENGYDRIATLHYQRREQDNDSIRRHEIDRGSEV
GA_CAD86C_Var5    ------------------------------------------------------------------
CAD86C_Ref        QSFRRDGPNIIYTKEMQLQEAFANKHKEYIEDLENGYDRIATLHYQRREQDNDSIRRHEIDRGSEV

cons

GA_CAD86C_Var1    GVFLKSDVDKNLHDKTDSKTKHEYSEKVRAPSTHGRDQYFIKEGNTEILRLVTRGKNEEERYVNLP
GA_CAD86C_Var2    GVFLKSDVDKNLHDKTDSKTKHEYSEKVRAPSTHGRDQYFIKEGNTEILRLVTRGKNEEERYVNLP
GA_CAD86C_Var3    ------------------------------------------------------------------
GA_CAD86C_Var4    GVFLKSDVDKNLHDKTDSKTKHEYSEKVRAPSTHGRDQYFIKEGNTEILRLVTRGKNEEERYVNLP
GA_CAD86C_Var5    ------------------------------------------------------------------
CAD86C_Ref        GVFLKSDVDKNLHDKTDSKTKHEYSEKVRAPSTHGRDQYFIKEGNTEILRLVTRGKNEEERYVNLP

cons

GA_CAD86C_Var1    AQQQRPVTLIPHTQYVVVDSGKNLLMERFIREQGEEAKNIRERMSKVTDLDNISNGKDAKSYGQRS
GA_CAD86C_Var2    AQQQRPVTLIPHTQYVVVDSGKNLLMERFIREQGEEAKNIRERMSKVTDLDNISNGKDAKSYGQRS
GA_CAD86C_Var3    ------------------------------------------------------------------
GA_CAD86C_Var4    AQQQRPVTLIPHTQYVVVDSGKNLLMERFIREQGEEAKNIRERMSKVTDLDNISNGKDAKSYGQRS
GA_CAD86C_Var5    ------------------------------------------------------------------
CAD86C_Ref        AQQQRPVTLIPHTQYVVVDSGKNLLMERFIREQGEEAKNIRERMSKVTDLDNISNGKDAKSYGQRS

cons

GA_CAD86C_Var1    QVGSEGQNQFHEYTNLHPEVPGTVPLKTDYLQSALLEMQNKSTIHQELLESSLRKQNELLHQILIE
GA_CAD86C_Var2    QVGSEGQNQFHEYTNLHPEVPGTVPLKTDYLQSALLEMQNKSTIHQELLESSLRKQNELLHQILIE
GA_CAD86C_Var3    ------------------------------------------------------------------
GA_CAD86C_Var4    QVGSEGQNQFHEYTNLHPEVPGTVPLKTDYLQSALLEMQNKSTIHQELLESSLRKQNELLHQILIE
GA_CAD86C_Var5    ------------------------------------------------------------------
CAD86C_Ref        QVGSEGQNQFHEYTNLHPEVPGTVPLKTDYLQSALLEMQNKSTIHQELLESSLRKQNELLHQILIE

cons

GA_CAD86C_Var1    RERMLQNQETASQVESKLETQSLPGHSVMATQTECHIGTQTEPQLMKPTRRKARSDNDSYSEDESQ
GA_CAD86C_Var2    RERMLQNQETASQVESKLETQSLPGHSVMATQTECHIGTQTEPQLMKPTRRKARSDNDSYSEDESQ
GA_CAD86C_Var3    ------------------------------------------------------------------
GA_CAD86C_Var4    RERMLQNQETASQVESKLETQSLPGHSVMATQTECHIGTQTEPQLMKPTRRKARSDNDSYSEDESQ
GA_CAD86C_Var5    ------------------------------------------------------------------
CAD86C_Ref        RERMLQNQETASQVESKLETQSLPGHSVMATQTECHIGTQTEPQLMKPTRRKARSDNDSYSEDESQ

cons

GA_CAD86C_Var1    MVVEDETKKVAWVKKRKPKKKLKHKDPRRSMRVYDLSRKIKTPILEESEVSPSLESEKHIKISKTN
GA_CAD86C_Var2    MVVEDETKKVAWVKKRKPKKKLKHKDPRRSMRVYDLSRKIKTPILEESEVSPSLESEKHIKISKTN
GA_CAD86C_Var3    ------------------------------------------------------------------
GA_CAD86C_Var4    MVVEDETKKVAWVKKRKPKKKLKHKDPRRSMRVYDLSRKIKTPILEESEVSPSLESEKHIKISKTN
GA_CAD86C_Var5    ------------------------------------------------------------------
CAD86C_Ref        MVVEDETKKVAWVKKRKPKKKLKHKDPRRSMRVYDLSRKIKTPILEESEVSPSLESEKHIKISKTN

cons

GA_CAD86C_Var1    EREEHIKNYGDMTRSTVTTSKNETVSSTQIKISDSDRKSRLRRDVLMEISDSIDDKVESDISDRRL
GA_CAD86C_Var2    EREEHIKNYGDMTRSTVTTSKNETVSSTQIKISDSDRKSRLRRDVLMEISDSIDDKVESDISDRRL
GA_CAD86C_Var3    ------------------------------------------------------------------
GA_CAD86C_Var4    EREEHIKNYGDMTRSTVTTSKNETVSSTQIKISDSDRKSRLRRDVLMEISDSIDDKVESDISDRRL
GA_CAD86C_Var5    ------------------------------------------------------------------
CAD86C_Ref        EREEHIKNYGDMTRSTVTTSKNETVSSTQIKISDSDRKSRLRRDVLMEISDSIDDKVESDISDRRL

cons

GA_CAD86C_Var1    RLQKQFSVTTDDTQLKSPLAIENDSNDNKSRSSSASKDHRNLVFSRQGSSTEAREIAAEHSSAKSK
GA_CAD86C_Var2    RLQKQFSVTTDDTQLKSPLAIENDSNDNKSRSSSASKDHRNLVFSRQGSSTEAREIAAEHSSAKSK
GA_CAD86C_Var3    ------------------------------------------------------------------
GA_CAD86C_Var4    RLQKQFSVTTDDTQLKSPLAIENDSNDNKSRSSSASKDHRNLVFSRQGSSTEAREIAAEHSSAKSK
GA_CAD86C_Var5    ------------------------------------------------------------------
CAD86C_Ref        RLQKQFSVTTDDTQLKSPLAIENDSNDNKSRSSSASKDHRNLVFSRQGSSTEAREIAAEHSSAKSK
```

```
cons


GA_CAD86C_Var1    SSDVEAQSSKLDSPSNPEPSKRVNEASESTKSSESVKPTQKSLPRYMQWYGKKSAAAKSTPAEKVV
GA_CAD86C_Var2    SSDVEAQSSKLDSPSNPEPSKRVNEASESTKSSESVKPTQKSLPRYMQWYGKKSAAAKSTPAEKVV
GA_CAD86C_Var3    ------------------------------------------------------------------
GA_CAD86C_Var4    SSDVEAQSSKLDSPSNPEPSKRVNEASESTKSSESVKPTQKSLPRYMQWYGKKSAAAKSTPAEKVV
GA_CAD86C_Var5    ------------------------------------------------------------------
CAD86C_Ref        SSDVEAQSSKLDSPSNPEPSKRVNEASESTKSSESVKPTQKSLPRYMQWYGKKSAAAKSTPAEKVV

cons


GA_CAD86C_Var1    PSKPKRSSKTKNDDDKVGRYGKIMNKDTQGDSEVKKNFKSKESEFIHPRILKEEKVTPVPEGPLPD
GA_CAD86C_Var2    PSKPKRSSKTKNDDDKVGRYGKIMNKDTQGDSEVKKNFKSKESEFIHPRILKEEKVTPVPEGPLPD
GA_CAD86C_Var3    ------------------------------------------------------------------
GA_CAD86C_Var4    PSKPKRSSKTKNDDDKVGRYGKIMNKDTQGDSEVKKNFKSKESEFIHPRILKEEKVTPVPEGPLPD
GA_CAD86C_Var5    ------------------------------------------------------------------
CAD86C_Ref        PSKPKRSSKTKNDDDKVGRYGKIMNKDTQGDSEVKKNFKSKESEFIHPRILKEEKVTPVPEGPLPD

cons


GA_CAD86C_Var1    VHPLLQHSEHRYEHQYENQNPLCYIQPTHIPKYLGQQPMPRKQAPEQQPIYVNQDDVKEPTQDIAE
GA_CAD86C_Var2    VHPLLQHSEHRYEHQYENQNPLCYIQPTHIPKYLGQQPMPRKQAPEQQPIYVNQDDVKEPTQDIAE
GA_CAD86C_Var3    ------------------------------------------------------------------
GA_CAD86C_Var4    VHPLLQHSEHRYEHQYENQNPLCYIQPTHIPKYLGQQPMPRKQAPEQQPIYVNQDDVKEPKQDIAE
GA_CAD86C_Var5    ------------------------------------------------------------------
CAD86C_Ref        VHPLLQHSEHRYEHQYENQNPLCYIQPTHIPKYLGQQPMPRKQAPEQQPIYVNQDDVKEPTQDIAE

cons


GA_CAD86C_Var1    SALTHSISISTSYEEDRKNPEVHVSKINIGGDNIPDIAHRSVTSKIDDNDSGIAMNTLVQQNNTKR
GA_CAD86C_Var2    SALTHSISISTSYEEDRKNPEVHVSKINIGGDNIPDIAHRSVTSKIDDNDSGIAMNTLVQQNNTKR
GA_CAD86C_Var3    ------------------------------------------------------------------
GA_CAD86C_Var4    SALTHSISISTSYEEDRKNPEVHVSKINIGGDNIPDIAHRSVTSKIDDNDSGIAMNTLVQQNNTKR
GA_CAD86C_Var5    ------------------------------------------------------------------
CAD86C_Ref        SALTHSISISTSYEEDRKNPEVHVSKINIGGDNIPDIAHRSVTSKIDDNDSGIAMNTLVQQNNTKR

cons


GA_CAD86C_Var1    LPITEKKSVFTIAYDDVQTKQLRPDSSSTSY
GA_CAD86C_Var2    LPITEKKSVFTIAYDDVQTKQLRPDSSSTSY
GA_CAD86C_Var3    -------------------------------
GA_CAD86C_Var4    LPITEKKSVFTIAYDDVQTKQLRPDSSSTSY
GA_CAD86C_Var5    -------------------------------
CAD86C_Ref        LPITEKKSVFTIAYDDVQTKQLRPDSSSTSY

cons
```
