## Supplementary material for "Mutations in a novel cadherin gene associated with Bt resistance in *Helicoverpa zea*": Supplemental_Figure_Captions.docx

**Supplemental Figure 1.** Distribution of the differences in nucleotide diversity values for permuted and resampled GA and GA-R populations at the first PCR amplified *cad-86C* locus (primer pair 1b; amplicon length = 421 bp).

**Supplemental Figure 2.** Distribution of the differences in nucleotide diversity values for permuted and resampled GA and GA-R populations at the second PCR amplified *cad-86C* locus (primer pair 2b; amplicon length = 520 bp).

**Supplemental Figure 3.** Full length cad-86C cDNA amplicons as detected by PCR and gel electrophoresis on a 1% agarose gel. Each well represents cDNA amplicons from the midgut of a single *H. zea* individual. Individuals 3-15 were from the GA-R line, while individuals 16-30 were from the GA line.

**Supplemental Figure 4.** Protein sequence alignment for each of the 5 variants detected in the GA and GA-R populations of *H. zea*.
