## Supplementary material for "Mutations in a novel cadherin gene associated with Bt resistance in *Helicoverpa zea*": Hzea_WGS_Supplemental_Methods.docx

*ExoAP Clean Protocol*

The ExoAP mastermix is made by mixing 5.3uL Exonuclease I (NEB product M0293S at 20,000 U/mL), 5.3uL Antarctic Phosphotase (NEB product M0289S at 5,000 U/mL) and 5.3 uL Antarctic Phosphatase Buffer (NEB product B0289S at 10x concentration). These were brought up to a total volume of 106 mL with PCR grade water. Once made, the mix can be added to PCR products in a ratio of 1 part ExoAP to 3 parts PCR product, and incubated in a thermal cycler under the following conditions: 37ºC for 15 min, 80ºC for 15 min, then held at 15ºC until Sanger sequencing submission.
