## Supplementary material for "Mutations in a novel cadherin gene associated with Bt resistance in *Helicoverpa zea*": Hzea_WGS_Supplemental_Tables1-3_and_5-7.docx

**Table S1.** Primer sequences for targeted portions of the *cad-86C* gene in *Helicoverpa zea.*

| **Primer Name** | **Forward Sequence** | **Reverse Sequence** |
| --- | --- | --- |
| Cad_1b | 5’-CAACTTGTGGTAACGGCTTTG | 5’-TGCTTATCCTCGTGTGAATTGT |
| Cad_2b | 5’-TCCTGATAAATCACAACTCCATCT | 5’-ACGGCAAAGAGGCATTCA |

**Table S2.** qPCR primer sequences for *cad-86C* and an endogenous control gene, α-Tubulin, in *Helicoverpa zea.*

| **Target Gene** | **Primer Name** | **Forward Sequence** | **Reverse Sequence** |
| --- | --- | --- | --- |
| Cadherin-86c | Cad_qPCR3 | 5’-CGGTAGTGATTCTGGTCTCT | 5’-GGAGGCATATAAAGTGGACATC |
| α-Tubulin | αTub_qPCR1 | 5’-CATGTTGTACCGTGGAGACG | 5’-CTGGTAGTTGATGCCCACCT |

**Table S3.** Barcoded forward primers and the reverse primer for *cad-86C* cDNA amplification.

| **Primer Name** | **Sequence** |
| --- | --- |
| cadUTR_F_bar1 | 5’-GTCATCACTAAACCAAGGGTGGAGTAGTATAACGGTAGCC |
| cadUTR_F_bar2 | 5’-GTCATCACTAACCTACCTGTGGAGTAGTATAACGGTAGCC |
| cadUTR_F_bar3 | 5’-GTCATCACTAACGTGTTGGTGGAGTAGTATAACGGTAGCC |
| cadUTR_F_bar4 | 5’-GTCATCACTAACTGGACTGTGGAGTAGTATAACGGTAGCC |
| cadUTR_F_bar5 | 5’-GTCATCACTAAGAGACTGGTGGAGTAGTATAACGGTAGCC |
| cadUTR_F_bar6 | 5’-GTCATCACTAAGTCGACTGTGGAGTAGTATAACGGTAGCC |
| cadUTR_F_bar7 | 5’-GTCATCACTAATATGCCGGTGGAGTAGTATAACGGTAGCC |
| cadUTR_F_bar8 | 5’-GTCATCACTACAACCATGGTGGAGTAGTATAACGGTAGCC |
| cadUTR_F_bar9 | 5’-GTCATCACTACACAGTGTGTGGAGTAGTATAACGGTAGCC |
| cadUTR_F_bar10 | 5’-GTCATCACTACAGAAGTGGTGGAGTAGTATAACGGTAGCC |
| cadUTR_F_bar11 | 5’-GTCATCACTACAGTGACTGTGGAGTAGTATAACGGTAGCC |
| cadUTR_F_bar12 | 5’-GTCATCACTACATGTGGTGTGGAGTAGTATAACGGTAGCC |
| cadUTR_F_bar13 | 5’-GTCATCACTACCTACCATGTGGAGTAGTATAACGGTAGCC |
| cadUTR_F_bar14 | 5’-GTCATCACTACGAACGTTGTGGAGTAGTATAACGGTAGCC |
| cadUTR_F_bar15 | 5’-GTCATCACTACGTAGGAAGTGGAGTAGTATAACGGTAGCC |
| cadUTR_F_bar16 | 5’-GTCATCACTACTAGTGGTGTGGAGTAGTATAACGGTAGCC |
| cadUTR_F_bar17 | 5’-GTCATCACTACTCTGACTGTGGAGTAGTATAACGGTAGCC |
| cadUTR_F_bar18 | 5’-GTCATCACTAGAACCTTGGTGGAGTAGTATAACGGTAGCC |
| cadUTR_F_bar19 | 5’-GTCATCACTAGACATCTGGTGGAGTAGTATAACGGTAGCC |
| cadUTR_F_bar20 | 5’-GTCATCACTAGAGAGACTGTGGAGTAGTATAACGGTAGCC |
| cadUTR_F_bar21 | 5’-GTCATCACTAGATCGAAGGTGGAGTAGTATAACGGTAGCC |
| cadUTR_F_bar22 | 5’-GTCATCACTAGCAATAGGGTGGAGTAGTATAACGGTAGCC |
| cadUTR_F_bar23 | 5’-GTCATCACTAGCTACCTTGTGGAGTAGTATAACGGTAGCC |
| cadUTR_F_bar24 | 5’-GTCATCACTAGGATATGGGTGGAGTAGTATAACGGTAGCC |
| cadUTR_F_bar25 | 5’-GTCATCACTAGTCATCAGGTGGAGTAGTATAACGGTAGCC |
| cadUTR_F_bar26 | 5’-GTCATCACTAGTGAGTCTGTGGAGTAGTATAACGGTAGCC |
| cadUTR_F_bar27 | 5’-GTCATCACTAGTTCGATGGTGGAGTAGTATAACGGTAGCC |
| cadUTR_F_bar28 | 5’-GTCATCACTATACGATCGGTGGAGTAGTATAACGGTAGCC |
| cadUTR_F_bar29 | 5’-GTCATCACTATCACTCTGGTGGAGTAGTATAACGGTAGCC |
| cadUTR_F_bar30 | 5’-GTCATCACTATCTCCAGTGTGGAGTAGTATAACGGTAGCC |
| cadUTR_F_bar31 | 5’-GTCATCACTATGACTCAGGTGGAGTAGTATAACGGTAGCC |
| cadUTR_F_bar32 | 5’-GTCATCACTATGGTTCCTGTGGAGTAGTATAACGGTAGCC |
| cadUTR_F_bar33 | 5’-GTCATCACTATGTGACTGGTGGAGTAGTATAACGGTAGCC |
| cadUTR_F_bar34 | 5’-GTCATCACTATTCGATGGGTGGAGTAGTATAACGGTAGCC |
| cadUTR_R1 | 5’-TCAATCGACGGACACGAATAC |

**Table S4.** Summary of Illumina reads collected for each individual. Shown are the total number of reads obtained for each individual, retained reads after filtering out low quality scores, total reads mapped to the *Helicoverpa zea* reference genome, and the mean sequencing depth.

| **Sample ID** | **Total PE Reads** | **Filtered PE Reads** | **Mapped Reads** | **Mean sequence depth (×)** |
| --- | --- | --- | --- | --- |
| B1 | 66,745,362 | 55,600,073 | 25,856,243 | 23.20 |
| C1 | 46,778,103 | 37,872,611 | 18,701,181 | 16.77 |
| D1 | 36,901,929 | 30,932,529 | 14,965,749 | 13.43 |
| D2 | 39,279,497 | 31,755,375 | 16,842,162 | 15.11 |
| D3 | 56,513,308 | 47,456,576 | 24,214,897 | 21.72 |
| E1 | 46,111,320 | 38,826,792 | 19,755,879 | 17.72 |
| E3 | 27,045,423 | 22,790,570 | 11,657,375 | 10.46 |
| F2 | 43,250,346 | 36,592,604 | 19,233,797 | 17.26 |
| G2 | 21,228,969 | 17,772,582 | 8,876,363 | 7.96 |
| H1 | 43,282,524 | 36,602,796 | 18,937,097 | 16.99 |

**Table S5.** See separate spreadsheet.

**Table S6.** Percentage identity matrix from T-coffee alignments (Noterdame et al. 2010) between *H. zea* CAD-86C protein (ID=537580) and five other cadherin proteins with known roles in Bt resistance.

|  | AAM78590.1 | AF367362.1 | AF519180 | AGG36450.2 | AY198374.1 | ID=537580 |
| --- | --- | --- | --- | --- | --- | --- |
| AAM78590.1 | 100 | 56 | 56 | 19 | 56 | 19 |
| AF367362.1 | 56 | 100 | 84 | 22 | 57 | 21 |
| AF519180 | 56 | 84 | 100 | 22 | 57 | 22 |
| AGG36450.2 | 19 | 22 | 22 | 100 | 21 | 17 |
| AY198374.1 | 56 | 57 | 57 | 21 | 100 | 21 |
| ID=537580 | 19 | 21 | 22 | 17 | 21 | 100 |

>AY198374.1 *Pectinophora gossypiella* cadherin-like protein (BtR)

>AF367362.1 *Heliothis virescens* cadherin-like protein mRNA

>AF519180 *Helicoverpa armigera* cadherin-like protein (Bt-Rm) mRNA

>AAM78590.1 midgut cadherin-like protein [*Chilo suppressalis*]

>AGG36450.2 cadherin-like protein 2 [*Chilo suppressalis*]

>ID=537580 *H. zea* CAD-86C

**Table S7.** PacBio reads number before and after length filtering, minimum length = 5000 bp. Individuals GA1 through GA15 were from the GA-R line, while individuals GA16 through GA30 were from the GA line.

| Samples | Before filtering | After filtering | Category |
| --- | --- | --- | --- |
| GA1 | 1 | 0 | Resistant |
| GA2 | 13 | 0 | Resistant |
| GA3 | 964 | 667 | Resistant |
| GA4 | 868 | 615 | Resistant |
| GA5 | 1006 | 697 | Resistant |
| GA6 | 948 | 653 | Resistant |
| GA7 | 0 | 0 | Resistant |
| GA8 | 4616 | 253 | Resistant |
| GA9 | 2391 | 383 | Resistant |
| GA10 | 2114 | 198 | Resistant |
| GA11 | 1552 | 344 | Resistant |
| GA12 | 4 | 2 | Resistant |
| GA13 | 10640 | 234 | Resistant |
| GA14 | 3990 | 180 | Resistant |
| GA15 | 1752 | 91 | Resistant |
| GA16 | 1822 | 1135 | Susceptible |
| GA17 | 2908 | 1947 | Susceptible |
| GA18 | 2722 | 1774 | Susceptible |
| GA19 | 2550 | 1697 | Susceptible |
| GA20 | 6 | 3 | Susceptible |
| GA21 | 1928 | 1272 | Susceptible |
| GA22 | 0 | 0 | Susceptible |
| GA23 | 0 | 0 | Susceptible |
| GA24 | 744 | 320 | Susceptible |
| GA25 | 2424 | 190 | Susceptible |
| GA26 | 1614 | 8 | Susceptible |
| GA27 | 2744 | 830 | Susceptible |
| GA28 | 1374 | 735 | Susceptible |
| GA29 | 3026 | 447 | Susceptible |
| GA30 | 2316 | 187 | Susceptible |
| Total | 57037 | 14862 |  |

**Table S8.** Homozygotes and heterozygotes for the five amino acid variants of *cad-86C*. GA-R individuals are denoted as GA3-15, and GA individuals are denoted as GA16-30.

| **Variants** | **Homozygotes** | **Heterozygotes** |
| --- | --- | --- |
| 1 | GA3, GA4, GA6, GA8, GA10, GA11, GA13, GA14, GA15, GA16, GA19, GA21, GA28, GA30 | GA5, GA27 |
| 2 |  | GA9 |
| 3 |  | GA9 |
| 4 | GA17, GA18, GA25, GA29 | GA5, GA27 |
| 5 | GA24 |  |
