## Supplementary figures and images for "Mutations in a novel cadherin gene associated with Bt resistance in *Helicoverpa zea*"

### FigS1_CAD86c1b_resamp_NucDivDiff_Dist.png

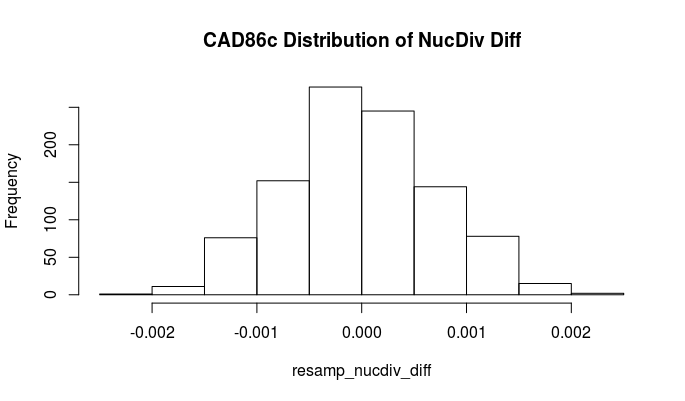

### FigS2_CAD86c2b_resamp_NucDivDiff_Dist.png

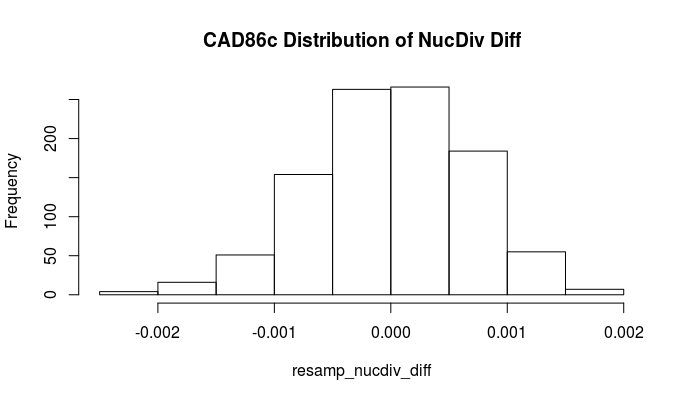

### FigureS3.jpg

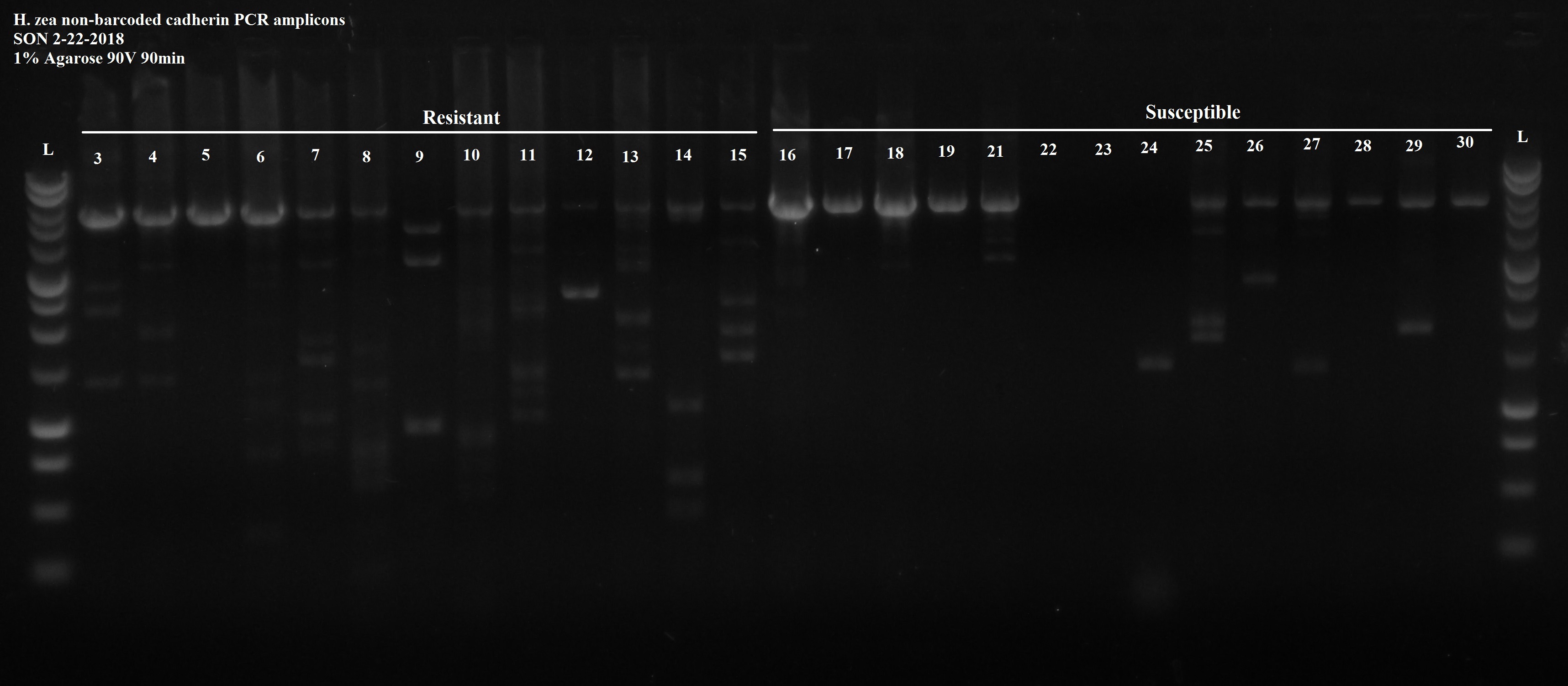
